# Reciprocity between retrograde signal and putative metalloprotease reconfigures plastidial metabolism and structure

**DOI:** 10.1101/2021.12.15.472781

**Authors:** Jin-Zheng Wang, Wilhelmina van de Ven, Yanmei Xiao, Xiang He, Haiyan Ke, Panyu Yang, Katayoon Dehesh

## Abstract

Reconfiguration of the plastidial proteome in response to environmental inputs is central to readjustment of its metabolic and structural states. This is necessary for the functionality of this metabolic hub, and the maintenance of organismal integrity. This report establishes the role of the plastidial retrograde signaling metabolite, MEcPP, in increasing the abundance of the putative plastidial metalloprotease (VIR3), and the ensuing decline of VIR3 target enzymes, ascorbate peroxidase and glyceraldehyde 3-phophate dehydrogenase B. The decreased abundance of these enzymes is linked to increased levels of their substrates: H_2_O_2_, an elicitor of salicylic acid production and stromule formation; and G3P the substrate for MEcPP synthesis. High-light treatment of wild type plants recapitulated the VIR3-associated reconfiguration of the plastidial metabolic and structural states. These results identify a previously unrecognized link between the stress-induced plastidial retrograde signaling metabolite and a putative zinc-binding metalloprotease. Moreover, the data reveal that the reciprocity between these two components, results in the reconfiguration of the metabolic and structural states of the plastid, deemed necessary to maintain cellular integrity and to shape adaptive responses.

## Introduction

Plastids are the metabolic, signaling and sensing centers well-equipped with a range of biochemical networks bound to maintain the quality and quantity of the plastidial proteome through regulated proteolysis and protein quality control mechanisms. To date, combined biochemical, genetic and proteomic studies have identified about 20 chloroplast protein-degrading machineries with more than 50 constituents (Nishimura *et al*, 2017). Among them a group of membrane bound proteases known as FtsH (**f**ilamentation **t**emperature **s**ensitive), initially identified in an *Escherichia coli* mutant with aberrant growth behavior (Santos & De Almeida, 1975). In plants FtsHs are strictly present in the endosymbiotic-derived organelles, mitochondria and chloroplasts. Hence, while *E. coli* contains only one FtsH, the *Arabidopsis* genome contains 17 *FtsH* sequences, twelve of which are known to code active enzymes and eight of which are exclusively targeted to chloroplasts (Ferro *et al*, 2010; Garcia-Lorenzo *et al*, 2006). Most these proteases contain an AAA (ATPase associated with various cellular activities) domain and a metalloprotease domain ligating a Zn^2+^ ion in the consensus sequence HEXXH (where X is any uncharged residue) (Sakamoto *et al*, 2003). In general, thylakoid localized FtsHs are critical in biogenesis of these membranes, and are also implicated in retrograde signaling. One such an example is FtsH2 (Var2), purposed to be involved in retrograde signaling via EXECUTER1 (EX1)-dependent mechanisms, whereby degradation of EX1 by FtsH2 is necessary for activation of the retrograde signaling pathway (Dogra *et al*, 2017; Wang *et al*, 2016).

Another member of metalloprotease superfamily is the stroma lamella localized VIRESCENT3 (VIR3) that lacks the ATP-binding domain but possesses the signature zinc-binding motif HEXXH at the C terminus (^235^HEAGH^239^) (Chen *et al*, 2000; Qi *et al*, 2016; Rawlings *et al*, 2014). The zinc-binding site is exposed to stroma and its perturbation by His^235^ substitution interferes with VIR3 function and/or stability (Qi *et al*., 2016). Loss of VIR3 function causes a virescent phenotype, hence the designation. It is suggested that VIR3 potentially exerts proteolytic activity as a monomer or in small complexes to maintain quality and quantity of the plastidial proteome (Qi *et al*., 2016).

Environmental perturbations result in increased production of reactive oxygen species (ROS) to lethal levels if unchecked. Ascorbate peroxidase (APX) is a multifamily of antioxidant isoenzymes originating from alternative splicing (Caverzan *et al*, 2012). The isoenzymes catalyze the conversion of H_2_O_2_ into H_2_O, under both standard and stress conditions in various subcellular compartments including cytosol, peroxisome, mitochondria, and chloroplast (stroma and membrane-bound thylakoid) (Kuo *et al*, 2020; Pandey *et al*, 2017; Shigeoka *et al*, 2002). Importantly, the thylakoidal isoform (tAPX) borders with the acceptor of photosystem I, and as such the first enzyme to intercept an H_2_O_2_ molecule produced (Huseynova *et al*, 2014).

Glucose metabolism is the foundation of all biochemical activities, and glyceraldehyde 3-phosphate dehydrogenase (GAPDH) is the ubiquitous multifamily enzyme comprised of glycolytic GAPDHs (GAPC), and highly regulated plastidial enzymes (GAPA and GAPB) catalyzing the reversible conversion of glyceraldehyde 3-phosphate (G3P) to 1, 3-bisphosphoglycerate (**1**,3BPG) with concomitant reduction of NAD(P)^+^ to NADH(P) in the photosynthetic reductive carbon cycle (Conley & Shih, 1995; Marri *et al*, 2005; Munoz-Bertomeu *et al*, 2009; Yang *et al*, 1993; Zaffagnini *et al*, 2013). In addition to their housekeeping roles, the GAPDHs are involved in non-traditional and diverse cellular functions such as influencing cell fate, regulation of ROS accumulation and cell death in response to pathogen attack (Colell *et al*, 2009; Henry *et al*, 2015). Indeed, one may further surmise expanded regulatory function of plastidial GAPDHs given that the first step of isoprenoids production by plastidial methylerythritol phosphate (MEP)-pathway is the condensation of pyruvate and G3P (C. Obiol-Pardo, 2011; Lange *et al*, 2000; Tritsch *et al*, 2010; Zeng & Dehesh, 2021).

Intriguingly, the MEP-pathway not only serves as a central plastidial biochemical route, but also as a sensing and signaling pathway confirmed by the functionality of its intermediate, 2-C-methyl-D-erythritol-2, 4-cyclopyrophosphate (MEcPP), as a precursor of isoprenoids and as a stress-specific retrograde signaling metabolite coordinating expression of selected stress-response nuclear genes (Benn *et al*, 2016; de Souza *et al*, 2017; Walley *et al*, 2015; Wang *et al*,2020; Xiao *et al*, 2012). The discovery of MEcPP as a retrograde signaling metabolite was founded on a genetic screen that led to the isolation of a mutant line designated *ceh1*, for constitutive expression of *Hydroperoxide Lyase* (*HPL*), an otherwise stress-inducible nuclear gene encoding a plastidial enzyme in the HPL branch of the oxylipin pathway (Xiao *et al*., 2012). This mutation is caused by a single amino-acid substitution in the penultimate MEP-pathway enzyme, hydroxymethylbutenyl diphosphate synthase (HDS), resulting in accumulation of MEcPP and the consequential growth retardation, in concert with high levels of salicylic acid (SA) and the induction of selected nuclear-encoded stress-response genes (Benn *et al*, 2014; Jiang *et al*, 2019; Wang *et al*, 2017; Xiao *et al*., 2012).

Towards identification of the genetic components of MEcPP signal transduction pathway, we performed a suppressor screen for revertants with fully or partially abolished *ceh1* phenotypes, in spite of high MEcPP levels. Here, we report reversion of selected *ceh1* mutant phenotypes by a mutant allele of *VIR3*. Next, we establish significance of zinc-binding motif in functional integrity of this putative metalloprotease, and further identify VIR3 interacting partners. Through biochemical, metabolomics and cell biological approaches we establish an inverse relationship between abundance of the enzyme and its interacting partners linked to a feedback loop for accumulation of MEcPP and by extension SA, and accompanied by formation of stromules, proposed conduits for information exchange between chloroplast and nucleus. Lastly, high-light stress recapitulation of the VIR3-associated multicomponent modification illustrate the biological relevance of these findings. In summary our results uncover a central component of the plastidial sensing, signaling and biochemical activities, reshaping the metabolic and structural states of this organelle in response to adverse environmental inputs.

## Results

### VIR3: a component of MEcPP signal transduction pathway

To identify the genetic components of the MEcPP-mediated signal transduction pathway, we mutagenized the *ceh1* mutant using ethyl methanesulfonate (EMS) and searched for revertants (*Rceh*) that despite high MEcPP levels display full or partial restoration of the *ceh1* phenotypes, such as reversion of dwarf stature, constitutive *HPL* expression, and high SA content. This led to identification of a mutant initially designated *Rceh4* (*Revertant of ceh1 line 4*), displaying partial recovery of *ceh1* dwarfed statue by ~50% (Fig. 1**a**, **b**). Targeted metabolomics analyses showed no reduction in the MEcPP levels, but a significant reduction (~70%) in SA, albeit with no alteration in high and constitutive *HPL*-driven luciferase activity, in the *Rceh4* compared to the *ceh1* mutant (Fig. 1**a**, **c**, **d**). Whole-genome sequencing identified *VIRESCENT3* (*VIR3*) as the candidate gene harboring single point mutation at the 4^th^ intron splicing site resulting in the protein frame shift (Fig. 1**e** and Supplemental Fig 1**a**-**b**). Bioinformatics studies identified the encoded protein as a member of metalloprotease superfamily lacking the ATP-binding domain but containing the signature zinc-binding motif HEXXH at the C terminus (^235^HEAGH^239^) (Fig. 1**e**) (Chen *et al*., 2000; Qi *et al*., 2016; Rawlings *et al*., 2014).

**Fig. 1.**
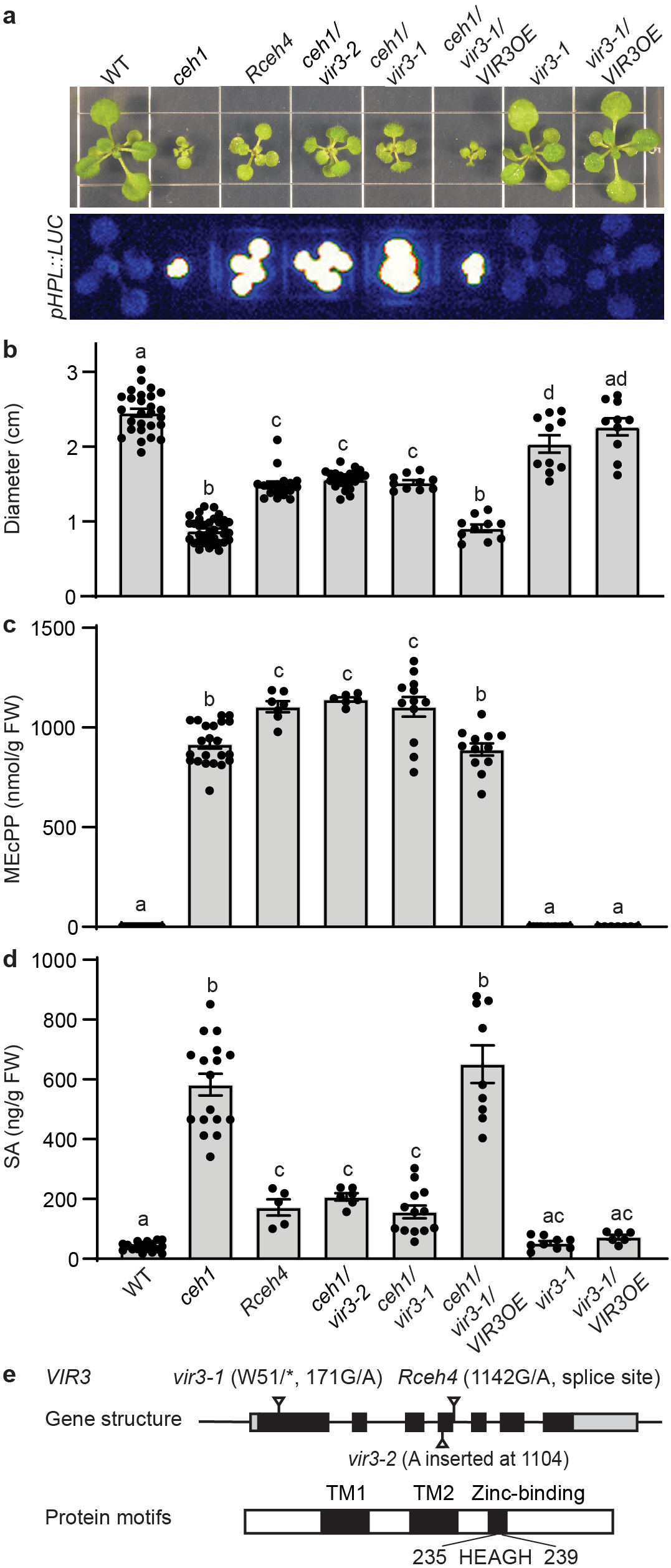
VIR3 is a component of MEcPP signal transduction pathway. (**a**) Representative images of 2 week-old WT, *ceh1*, *Rceh4*, *ceh1/vir3-1*, *ceh1/vir3-2*, *ceh1/vir3-1/VIR3OE*, *vir3-1*, and *vir3-1/VIR3OE* seedlings grown in long days (16h light/8h dark) (upper panel), and the corresponding dark-field images displaying *pHPL::LUC* activity (lower panel). (**b**) Seedling dimeter, (**c**) MEcPP and (**d**) SA levels of the aforementioned genotypes. Data (**b-d**) are mean ± SEM for each genotype with at least three biological replicates. Letters above bars indicate significant differences determined by one-way ANOVA with Tukey’s multiple comparisons test (P < 0.05). (**e**) Gene and protein structures of *VIR3*. Black boxes represent exons, lines represent introns and UTRs, and nucleotide and amino acid changes are displayed on the top or at the bottom of the top bar. The VIR3 transmembrane domains (TM) and the conserved zinc-binding domain (HEAGH) are shown on the lower panel bar.

To confirm *VIR3* as the gene responsible for partial reversion of *ceh1* aberrant phenotypes, we crossed *ceh1* to *vir3-1* loss of function mutant (*ceh1/vir3-1*) (Fig. 1**e**), In addition, we generated *ceh1* lines expressing <u>C</u>lustered <u>R</u>egularly <u>I</u>nterspaced <u>S</u>hort <u>P</u>alindromic <u>R</u>epeats (CRISPR)-based construct with guide RNA ~30bp upstream of *Rceh4* point mutation (herein designated *ceh1/vir3-2*). The *ceh1/vir3-2* expresses a premature VIR3 protein as the result of a nucleotide insertion at the 4^th^ exon (Fig. 1**e** and Supple Fig. 1**b**-**d**). The phenotypic and metabolomics analyses of *ceh1* backgrounds introgressed into two independent *vir3* loss of function mutants (*ceh1/vir3-1* and *ceh1/vir3-2*) matched those of the *Rceh4*, thereby confirming *vir3* as the mutant allele responsible for increasing the size and deceasing SA levels in the revertants, albeit with no impact on the constitutive expression of *HPL* (Fig. 1**a**-**e**). We further verified the finding by integrating *vir3-1/p35::VIR3-FLAG* (*vir3-1/VIR3OE*) (a gift from Prof. Fei Yu) into the *ceh1* mutant backgrounds (*ceh1/vir3-1/VIR3OE*). Detailed analyses of the genotype clearly show that overexpression of *VIR3* reverts *ceh1/vir3-1* partial size and SA level recoveries to those of the *ceh1* mutant, further verifying *Vir3* as the responsible allele (Fig. 1**a**-**e**). It is noteworthy that all the analyses indicate that under the standard growth condition *vir3-1* and *vir3-1/VIR3OE* lines are indistinguishable from each other and from the WT plants, supporting the functional association of VIR3 with stress, as evidenced by VIR3 functional input in the high MEcPP accumulating *ceh1* line.

To assess the potential interference of constitutively high MEcPP levels with the signal transduction pathway components, we further analyzed the DEX-inducible MEcPP accumulating genotypes in the WT type (*HDSi*) and *vir3* mutant backgrounds (*HDSi/vir3-1*). The result show that at 72h-post DEX induction seedlings display equally chlorotic phenotypes, together with reduced expression of *HDS* and in concert with enhanced MEcPP levels in both genotypes (Fig. 2**a**-**c**). In contrast however, DEX-induced MEcPP-mediated increase in SA levels remained restricted to *HDSi* line, and not detectable in the *HDSi/vir3-1* (Fig. 2**d**). This data confirms VIR3 as a component of MEcPP signal transduction pathway, with selective functional input in SA production.

**Fig. 2.**
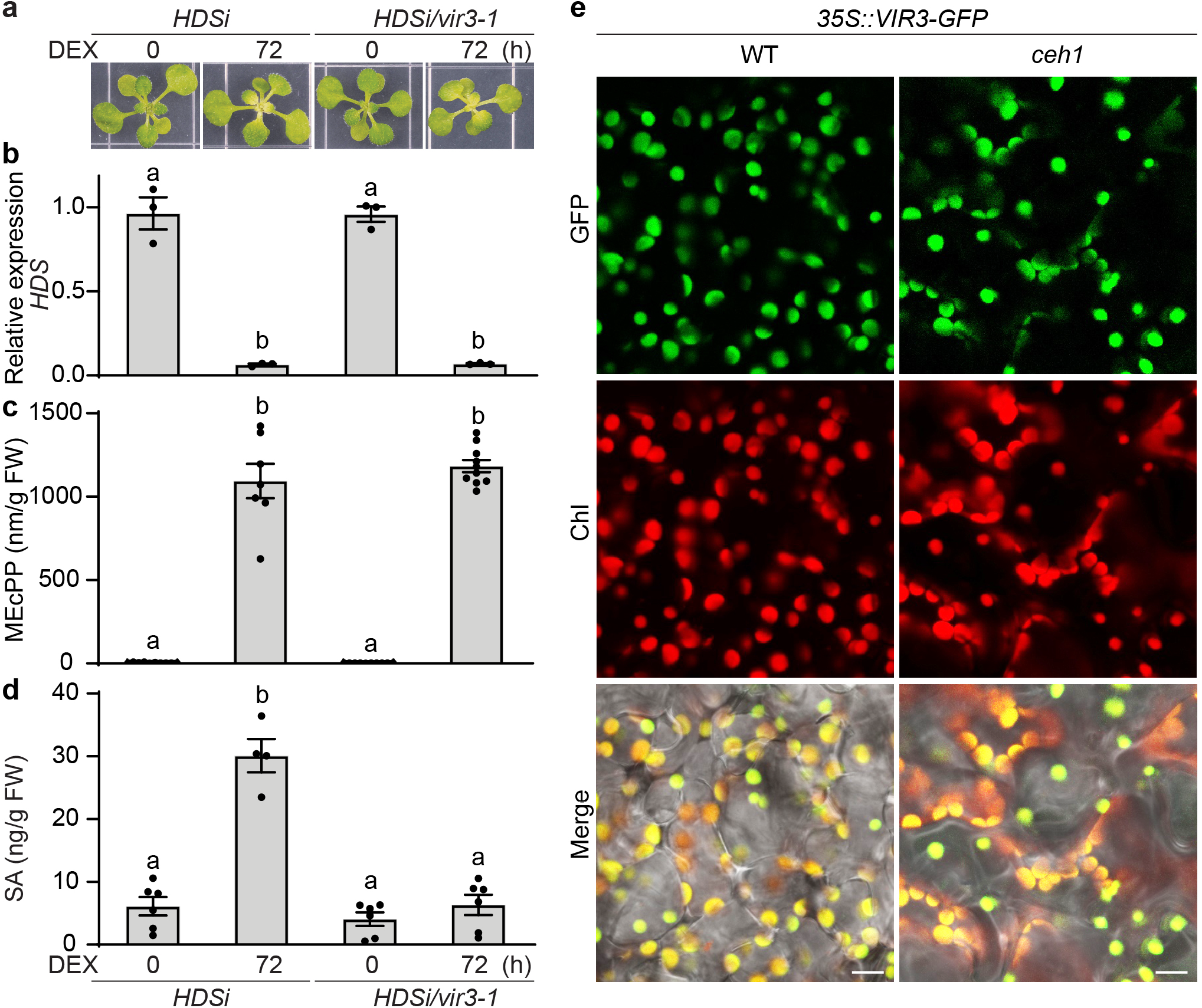
VIR3 is essential for MEcPP-mediated SA accumulation. (**a**) Representative images of 2-week-old untreated (0) or DEX-treated (post 72 h) *HDSi* and *HDSi/vir3-1* seedlings grown under long day conditions. (**b**) Relative expression levels of *HDS* in untreated (0) or DEX-treated (post 72 h) genotypes. (**c-d**) Levels of MEcPP (**c**) and SA (**d**) in the aforementioned genotypes. Data (**b-d**) are mean ± SEM for each genotype with at least three biological replicates. Statistical analyses were performed with one-way ANOVAwith Tukey’s multiple comparisons test (*P* < 0.05). (**e**) Plastidial localization of VIR3 in WT and *ceh1*. Bar=5μm.

To confirm plastidial localization of VIR3, and to exclude the potentials of its mislocalization as a consequence of high MEcPP levels in the *ceh1* mutant background, we used confocal microscopy and compared VIR3 localization in the WT versus the mutant by imaging the VIR3-GFP fusion in these plants (Fig. 2**e**). The data clearly verify the previously reported plastidial localization of VIR3 (Qi *et al*., 2016), and further confirm its unperturbed localization by high MEcPP content in the *ceh1* mutant.

### VIR3 function depends on the integrity of the zinc-binding motif

The presence of the zinc binding signature at the C terminus (^235^HEAGH^239^) of VIR3 led us to question the functional significance of this motif. Towards this goal we generated transgenic *vir3-1* lines expressing VIR3-FLAG and VIR3^H239Y^-FLAG constructs driven by UBQ10 promoter. We specifically exploited UBQ10-deriven expression construct of VIR3, since the weak expression levels driven by the native promoter proved prohibitive to the studies. Next, we examined the phenotypic characteristics of these genotypes along with the corresponding controls, namely WT, *vir3*, *vir3-1/VIR3OE*, under standard and constant light conditions. In contrast to the indistinguishable phenotypic characteristics of all the examined genotypes grown under standard conditions, the *vir3-1* and *vir3-1/pUBQ::VIR3^H239Y^-FLAG* genotypes displayed distinct variegation patterns not observed in the remainder lines grown under continuous light conditions (Fig. 3**a**).

**Fig. 3.**
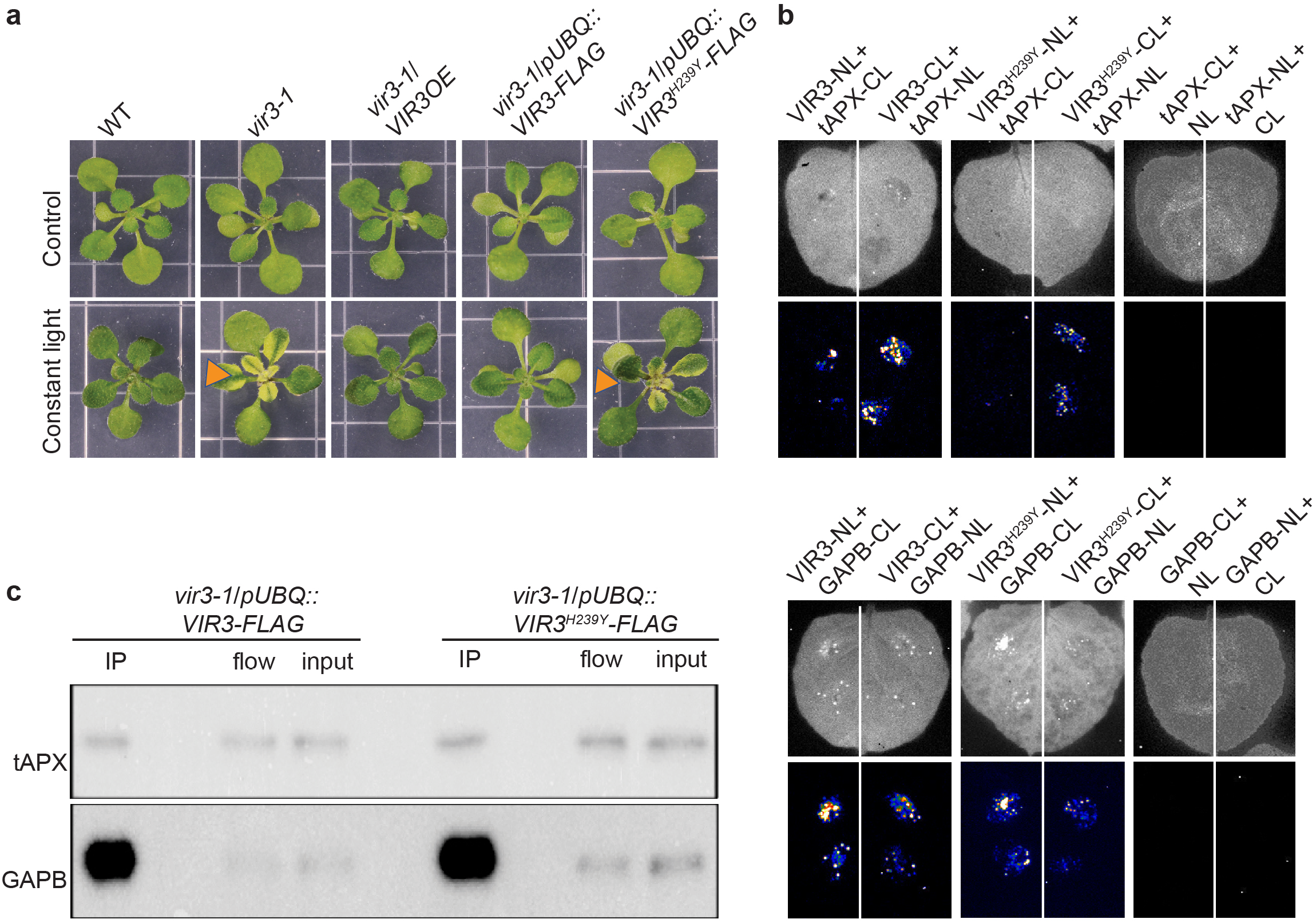
Zinc-binding motif is required for the functionality but not for VIR3 binding to tAPX and GAPB. (**a**) Representative images of WT, *vir3-1*, *vir3-1/VIR3OE*, *vir3-1/pUBQ::VIR3-FLAG* and *vir3-1/pUBQ::VIR3^H239Y^-FLAG* seedlings grown in long days (16h light/8h dark), without (control) or with exposure to 7 days of constant light leading to the exclusive development of variegation patterns on *vir3-1* and *vir3-1/pUBQ::VIR3^H239Y^-FLAG* leaves, but not in the other genotypes. (**b**) Representative images of infiltrated N. benthamiana leaves (top), and the corresponding split luciferase complementation assays (bottom), expressing permutations of VIR3 or VIR3^H239Y^ with tAPX (upper panel) or GAPB (lower panel), and each fused to N- and C-terminal fragments of luciferase (NL and CL respectively). Control consisted of LUC fusion constructs between tAPX and GAPB. LUC luminescence was analyzed after 2 days. The experiment were repeated at least three times. (**c**) Co-im-munoprecipitation (Co-IP) assay displays the in vivo interaction between VIR3 with tAPX and GAPB in *vir3-1/pUBQ::-VIR3-FLAG* and *vir3-1/pUBQ::VIR3^H239T^-FLAG* seedlings, using FLAG antibody for Co-IP, and tAPX and GAPB specific antibodies for immunoblot assays. The lanes between IP and flow are blank.

The data illustrate the significance of zinc-binding domain for VIR3 functionality, and further denote the stress-associated function of the enzyme.

### VIR3 physically interacts with tAPX and GAPB

The significance of zinc-binding domain in VIR3 functionality, strengthened the notion of its putative function as a metalloprotease. It is of note that the nature of VIR3 as a membrane bound protein hampered our efforts to examine the protease activity of recombinant VIR3 enzyme in an *in vitro* activity assay. However, the functionality of zinc-binding domain led us to question the existence of VIR3 interactive/substrate proteins. Towards this goal, we used *Nicotiana benthamiana* leaves transiently expressing *pUBQ::VIR3-FLAG* and *pUBQ::VIR3^H239Y^-FLAG* to immunoprecipitate VIR3 interactive proteins using FLAG antibody, followed by LC/MS analyses of the candidate proteins. Both constructs led to immunoprecipitation of several common proteins among then the two we initially focused on: the thylakoidal isoform of ascorbate peroxidase (tAPX; At1G77490) and plastidial GAPDH isoform B (GAPB; At1G42970) (Supplemental Table 1). This focus is based on the function of tAPX as an H_2_O_2_ scavenger, and the action of GAPB in the reversible conversion of the MEP-pathway initiating substrate, G3P to **1**,3BPG.

Next, we exploited split luciferase complementation assays in tobacco leaves transiently expressing different combinations of native and mutated *VIR3/tAPX* and *VIR3/GAPB* constructs fused to N- or C-terminal fragment of *luciferase* (*LUC*). To eliminate possible auto-luminescence, we checked LUC activity in leaves expressing *tAPX* and *GAPB* constructs fused to *LUC* (Fig. 3**b**). The Bioluminescence data clearly shows reconstitution of LUC activity in leaves co-infiltrated with VIR3 or VIR3^H239Y^ together with tAPX or GAPB, but not in the control leaves (Fig. 3**b** and Supplemental Fig. 2**a**). Specifically, both native VIR3 and VIR3^H239Y^ display notable physical interaction with tAPX and GAPB, with the strongest signals observed in leaves expressing VIR3-CL/ tAPX-NL, and in VIR3-NL/GAPB-CL expressing leaves.

To verify these interactions, we employed Arabidopsis *vir3-1* lines expressing VIR3-FLAG and VIR3^H239Y^-FLAG and performed co-immunoprecipitation (CO-IP) assay using a FLAG antibody, followed by immunoblot analyses using the tAPX and GAPB specific antibodies (Fig. 3**c** and Supplemental Fig. 2**b-c**). To examine instability of VIR3 previously observed in transgenic lines expressing VIR3^H235L^-FLAG zinc-binding domain (Qi *et al*., 2016), we first performed immunoblot analyses using FLAG antibody (Supplemental Fig. 2**b).**The presence of VIR3 in both transgenic lines exclude the possibility of protein instability as the result of VIR3 mutation. Next, we performed immunoblot analyses using tAPX and GAPB specific antibodies. The clear and specific presence of a tAPX and a GAPB reacting band in the IP fraction verified the *in vivo* interaction of VIR3 with the tAPX and GAPB proteins in stably transformed Arabidopsis lines (Fig. 3**c** and Supplemental Fig. 2**c**). As an additional control for excluding non-specific binding, we performed the immunoblots using tAPX and GAPB specific antibodies on beads incubated with protein extracts from nontransgenic wild type seedlings (supplemental Fig. 2**d**). Collectively, our data show that VIR3 physically interacts with tAPX and GAPB, and further excludes the significance of the integrity of zinc-binding domain for the interaction, while confirms significance of the zinc-binding domain for VIR3 functionality.

### MEcPP-mediated increase in VIR3 is inversely correlated with tAPX and GAPB abundance

Partial and selective suppression of *ceh1* phenotypes in *Rceh4*, *ceh1/vir3-1* and *ceh1/vir3-2* lines (Fig. 1**a**-**e**) prompted the question of potential impact of MEcPP accumulation on transcriptional and/or translational regulation of VIR3. To address this, we first compared VIR3 transcript levels in WT, *ceh1*, *vir3-1/VIR3OE and ceh1/ vir3-1/VIR3OE* lines (Fig. 4**a**). The presence of similar VIR3 transcript levels in *ceh1* versus the WT, and between *vir3-1/VIR3OE* and *ceh1/ vir3-1/VIR3OE* lines is a clear indication of unaltered expression levels as the result of high MEcPP accumulation. However, immunoblot analyses of FLAG tagged lines clearly showed enhanced VIR3 abundance in *ceh1/vir3-1/VIR3OE* compared to the levels in *vir3-1/VIR3OE* lines (Fig. 4**b**, and Supplemental Fig. 3**a-b**). Additional immunoblots analyses using FLAG tagged *HDSi/VIR3OE* lines before and 72h post DEX-induction confirmed the positive correlation between increased MEcPP accumulation and enhanced VIR3 abundance (Fig. 4**c** and Supplemental Fig. 3**c-d**).

**Fig. 4.**
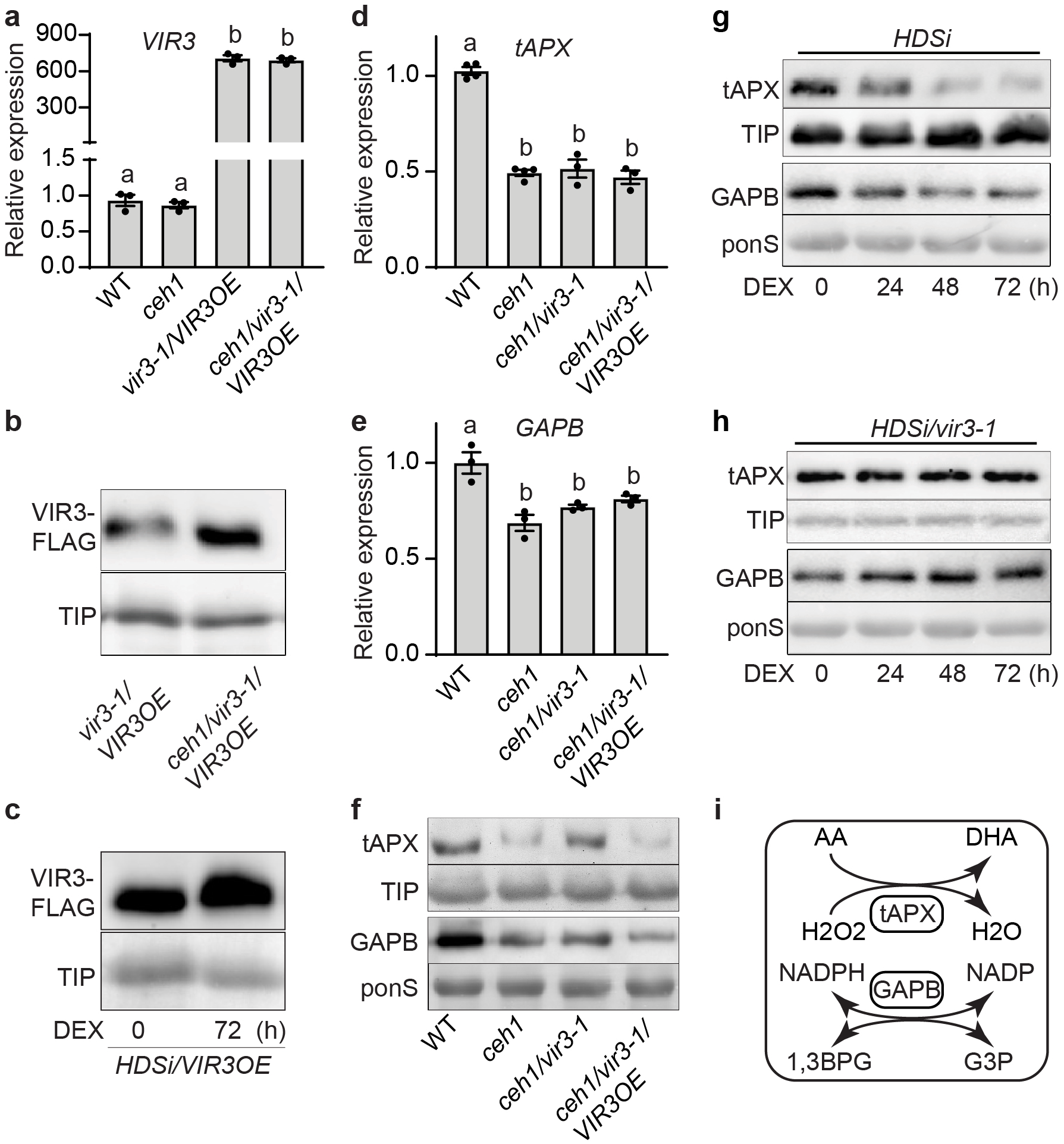
MEcPP-mediated increase in VIR3 is inversely correlated with tAPX and GAPB abundance. (**a**) Transcript levels of *VIR3* in WT, *ceh1*, *vir3-1/VIR3OE* and *ceh1/vir3-1/VIR3OE* seedlings, (b-c) Immunoblot analyses of VIR3 abundance using FLAG antibody applied on blot of protein extracts from *vir3-1/VIR3OE* and *ceh1/vir3-1/VIR3OE* (**b**), and (**c**) from untreated (0) and DEX-treated (72h) *HDSi/VIR3OE* seedlings, support MEcPP-mediated increase in VIR3 protein abundance. Specific antibody against TIP was used on the same blot to determine equal loading of the membrane bound proteins. (**d-e**) Transcript levels of tAPX (**d**), and GAPB (**e**) in WT, *ceh1*, *vir3-1/-VIR3OE* and *ceh1/vir3-1/VIR3OE*. Total RNAs isolated from 2-week-old seedlings grown under long-day (LD; 16 h light/8 h dark) condition was subjected to qRT-PCR analyses, and subsequently normalized to the levels of At4g264IO (M3E9). Data are mean ± SEM for each genotype with at least three biological replicates. Statistical analyses were performed with one-way ANOVA with Tukey’s multiple comparisons test, different letters indicate significant differences (*P* < 0.05). (f) Immunoblot analyses showing tAPX (upper panel) and GAPB (lower panel) protein abundance in WT, *ceh1*, *ceh1/vir3-1* and *ceh1/vir3-1/VIR3OE*. Specific antibody against TIP was used on the tAPX blot to show equal loading of the membrane bound protein, and Ponceau staining (ponS) was used for the loading control of the GABP, as the soluble protein. (g-h) Immunoblot analyses displaying tAPX and GAPB abundance in *HDSi* (g) and *HDSi/vir3-1* genotypes (h), at various hours post DEX-treatment (0, 24, 48 and 72 hours), using TIP and ponS as loading controls respectively. (i) Schematic presentation of tAPX function in conversion of H_2_O_2_ to water along with conversion of ascorbic acid (AA) to dehydroascorbic acid (DHA); and GAPB catalyzing the reversible conversion of glyceraldehyde 3-phosphate (G3P) to 1, 3-bisphosphoglycerate (1,3BPG) with concomitant reduction of NAD(P)^+^ to NADH(P).

Next we examined transcriptional and translational profiles of tAPX and GABP in WT, *ceh1*, *ceh1/ vir3-1 and ceh1/ vir3-1/VIR3OE* lines (Fig 4**d**-**f** and Supplemental Fig. 3**e**-**g**). This data shows reduced levels of *tAPX* and *GABP* expression in all the constitutively MEcPP accumulating *ceh1* backgrounds compared to the WT (Figs. 4**d** and **e**, and Supplemental Figs. 3**f** and **g**). In fact, this reduction in all the *ceh1* backgrounds is independent of the presence or absence of VIR3. In contrast, the tAPX and GABP proteins levels appear to be independent of the *ceh1* mutant background, but dependent on the presence of VIR3 (Fig. 4**f** and Supplemental Fig. 3**e**). This data specifically show a decrease in the abundance of both tAPX and GAPB proteins in *ceh1* and *ceh1/vir3-1/VIROE* compared to the WT and *ceh1/vir3-1*, thereby identifying VIR3 as a potential metalloprotease targeting these binding partners, albeit at different degrees.

Given the inverse relationship between the abundance of VIR3 and the levels of its binding proteins in the *ceh1* mutant background, we further examined the validity of this finding by testing tAPX and GAPB protein levels in the inducible MEcPP producing lines, *HDSi* and *HDSi/vir3-1* (Fig. 4**g**-**h** and Supplemental Fig. 3**h**-**m**). This data clearly shows reduced *tAPX* and *GAPB* protein levels post DEX-induction in *HDSi* but not in *HDSi/vir3* lines, thereby providing additional support to the notion of VIR3 function as a metalloprotease responsible for the degradation of these binding partners/substrates.

Collectively, the data provide experimental evidence for the likely MEcPP-mediated enhanced abundance/stability of VIR3 protein, but not the corresponding transcript levels in the *ceh1* mutant background, and further illustrate an inverse correlation between the abundance of VIR3 and its binding proteins, tAPX and GAPB.

### VIR3 functional input in ROS production and stromule formation

Given the tAPX function in intercepting H_2_O_2_ accumulation (Fig. 4**i**), we tested the consequential impact of reduced tAPX levels in *ceh1* compared to its seemingly stable levels in other genotypes (WT, *ceh1/vir3-1*, and *vir3-1*), using 3,3’-Diaminobenzidine (DAB) staining method (Fig. 5**a** and Supplemental Fig.4**a**). The representative images showing the extent of DAB staining in the various seedlings are well aligned with differential abundance of tAPX in these lines, specifically confirming the higher H_2_O_2_ levels in *ceh1* compared to the other genotypes.

**Fig. 5.**
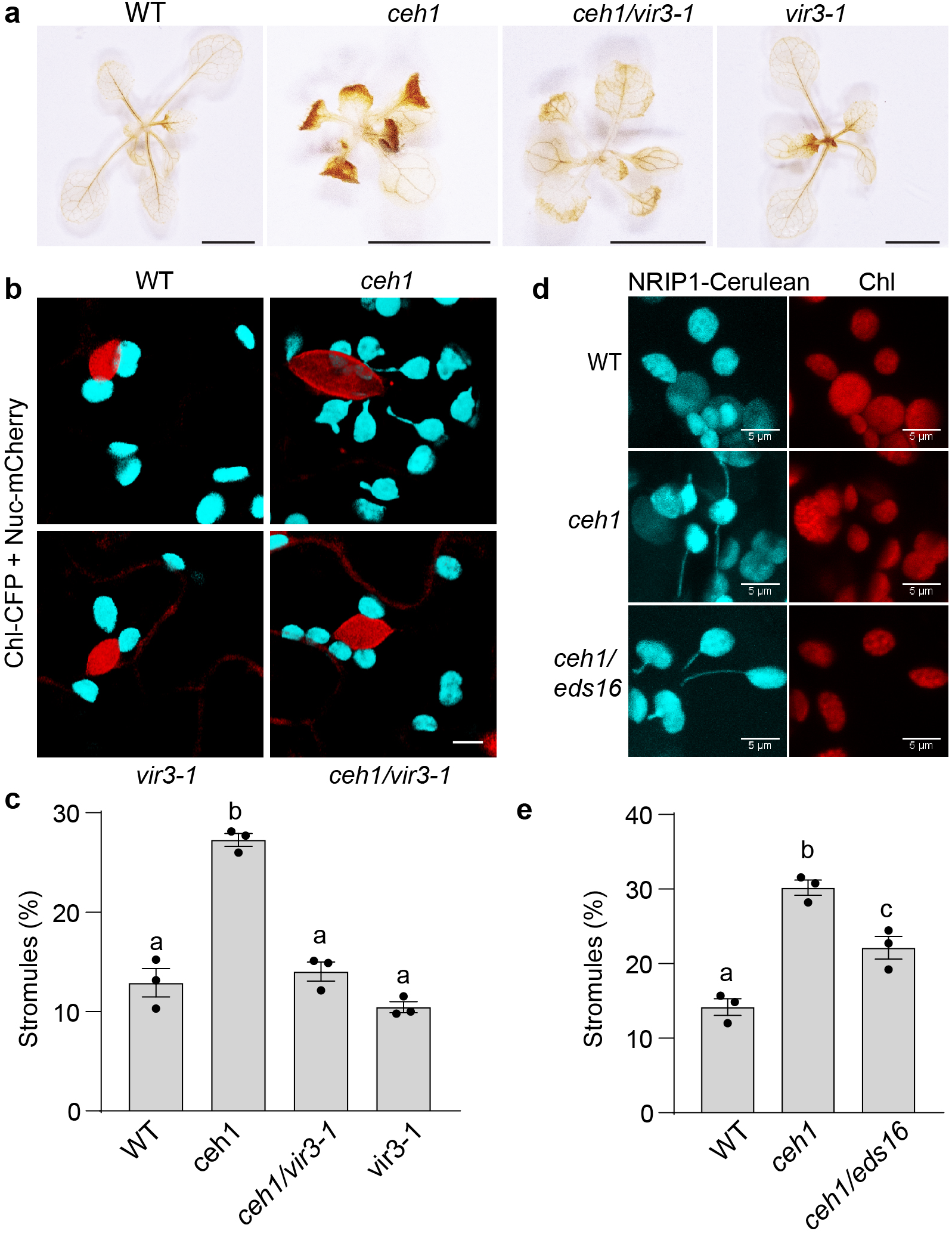
VIR3 functional input in ROS production and stromules formation. (**a**) Representative images of diaminobenzidine (DAB) stained 2-week-old WT, *ceh1*, *ceh1/vir3-1* and *vir3-1* seedlings show differential H_2_O_2_ accumulation in the genotypes. Bar=0.5cm. (**b-e**) Confocal images display nucleus (red) and chloroplast (cyan) without and with stromules in WT, *ceh1*, *ceh1/vir3-1* and *vir3-1* (**b**), and (**d**) WT, *ceh1* and *ceh1/eds16* seedlings with the images of chloroplast fluorescence, and their corresponding statistical analyses (**c** and **e**). The markers are CFP and mCherry fused to chloroplast transient peptide and to WPP nuclear localization signal, respectively. Bar=5μm. Data are mean ± SEM (n ≥ 3) for each genotype repeated three times with similar results. Statistical analyses were performed by one-way ANOVA with Tukey’s multiple comparisons test, (*P* < 0.05).

One of the reported functions of H_2_O_2_ is the induction of stroma-filled dynamic tubules structures that form in plastids, proposed to function as a conduit for transfer of information to nucleus (Caplan *et al*, 2015; Natesan *et al*, 2005). The differential levels of this signaling molecule prompted us to compare stromules formation in WT, *ceh1*, *ceh1*/*vir3-1* and *vir3-1* genotypes transformed with a construct simultaneously expressing chloroplast-targeted CFP and nuclear-targeted mCherry (Fig. 5**b-c** Supplemental Fig.4**b**). The confocal images followed by statistical analyses clearly portray higher % of stromules formed in *ceh1* compared to the other three genotypes examined. Moreover, the statistically significantly higher % of stromules, in conjunction with increased SA levels in *ceh1/vir3-1* compared to WT and *vir3-1* genotypes (Figs. 1**d** and 4**b**-**c**), led us to examine the potential contribution of SA to the formation of these conduits. Thus, we compared % of stromules formed in WT versus *ceh1*, and the SA deficient *ceh1* double mutant line (*ceh1/eds16-1*) lacking the SA-biosynthesis gene (Xiao *et al*., 2012) (Fig. 5**d**-**e** and Supplemental Fig.4**c**). The data is based on two independent lines each transformed with a different maker, namely NRIP1-Cerulean(Caplan *et al*., 2015) and RB-GFP. Both lines show significantly lower % of stromules in *ceh1/eds16* relative to the *ceh1*, but higher than the WT, establishing the previously reported contribution of SA to stromule formation (Caplan *et al*., 2015).

The result collectively provides a direct link between the lowered tAPX enzyme abundance and the consequential higher H_2_O_2_ levels to increased number of stromules in the *ceh1* mutant. Furthermore, the finding further establishes additive contributions of H_2_O_2_ and SA to the formation of these plastidial structures in the *ceh1* mutant backgrounds.

### High-light recapitulates VIR3-medited reconfiguration of the plastidial metabolic and structural states

To examine VIR3 functional role in plant adaptive responses, we exploited high-light (HL), a naturally occurring prevalent daily challenge. Next, we showed HL-mediated increase in MEcPP levels in WT and *vir3-1* mutant compared to the untreated lines (Fig. 6**a**). The notably higher MEcPP levels in the HL treated WT versus that of *vir3-1* alludes to a direct or an indirect VIR3 functional input in the MEP-pathway manifested by MEcPP levels. Next, we examined the correlation between HL-mediated increase of MEcPP and the potential enhanced abundance of VIR3 as previously observed in MEcPP accumulating *ceh1* and *HDSi* lines (Figs. 4**b** and **c**, and supplemental Fig. 3**a** and **c**). Because of the unavailability of a VIR3 specific antibody, and undetectable VIR3 levels in plants expressing VIR3-FLAG under the native promoter, HL assays were performed with *vir3-1/VIROE-FLAG* lines (Fig. 6**b** and Supplemental Fig. 5**a-b**). The data clearly show that even the overexpressing line exhibit higher levels of VIR3 in HL-treated versus untreated lines.

**Fig. 6.**
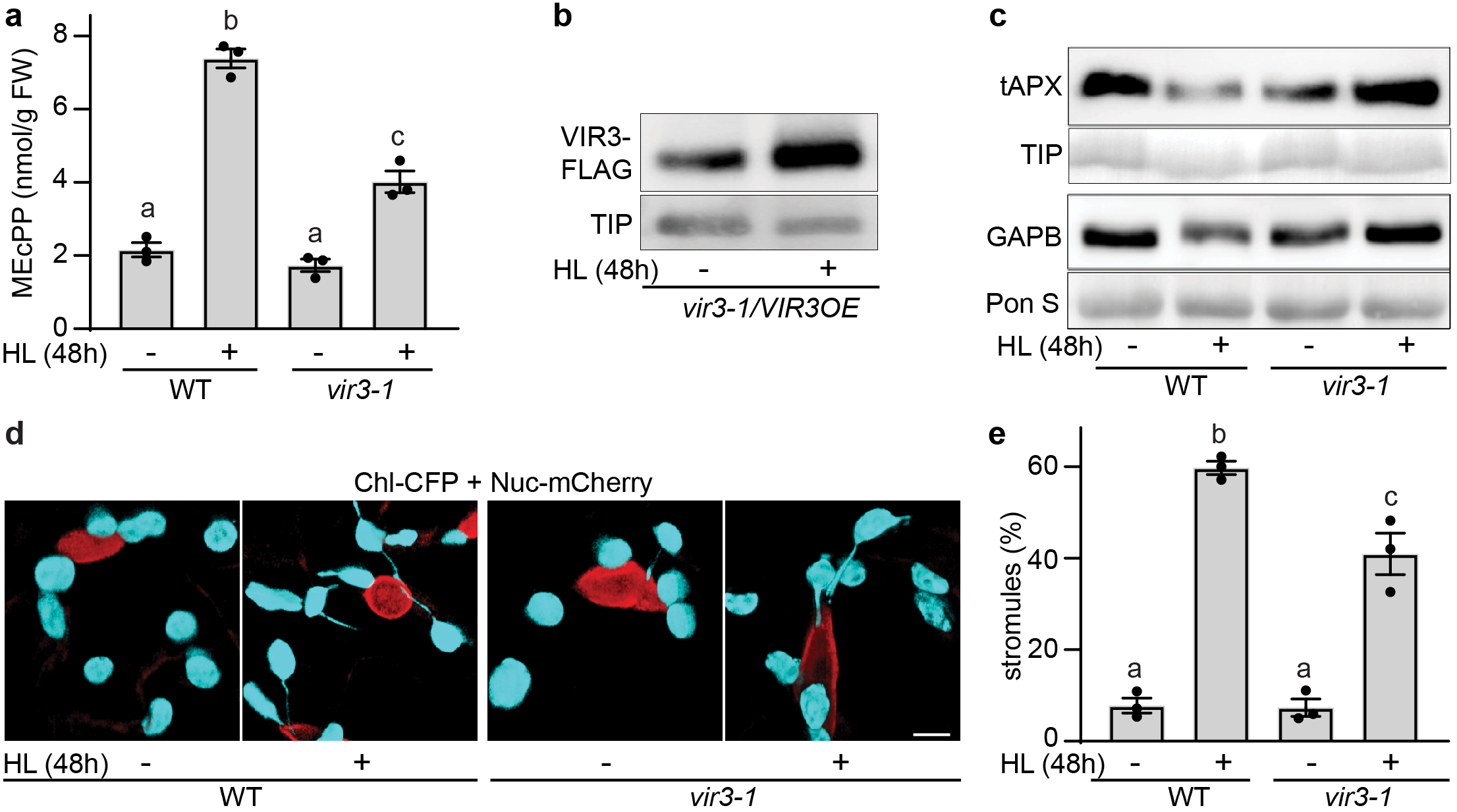
High-light reaffirms VIR3 role in reconfiguration of the plastidial metabolic and structural states. (**a**) Light induction of MEcPP accumulation in untreated (-) and 48 h high-light (HL)-treated (+) WT and *vir3-1* seedlings. (**b**) Immunoblot analyses of VIR3 abundance using FLAG antibody in untreated (-) and 48 h high-light (HL)-treated (+) *vir3-1/VIR3OE* seedlings. TIP antibody was used on the same blot as the loading control. (**c**) Immunoblot analyses of tAPX and GAPB protein levels in untreated (-) and 48h HL-treated (+) WT and *vir3-1* seedlings, using TIP antibody and ponceau staining (ponS) as loading controls respectively. (**d**) Representative confocal images showing nucleus (red) and chloroplasts (cyan) without and with stromules in untreated (-) and 48 h HL-treated (+) WT and *vir3-1* seedlings, and (e) their corresponding statistical analyses. Bar=5μm. Data are mean ± SEM (n ≥ 3). The experiment was repeated three times for each genotype. Statistical analyses were performed by one-way ANOVA with Tukey’s multiple comparisons test (*P* < 0.05).

Next, we compared the abundance of tAPX and GAPB in WT and *vir3-1* plants subjected to no or 48h of HL treatment (Fig. 6**c** and Supplemental Fig. 5**c-e**). The levels of both enzymes are reduced in the HL treated WT seedlings, but are increased in *vir3-1* mutant plant, supporting the inverse correlation between abundance of VIR3 and the levels of its interacting proteins.

Furthermore, confocal examination of stromule formation in untreated and HL-treated WT and *vir3-1* seedlings clearly illustrate presence of higher % of these dynamic strucutres in HL-treated seedlings, albeit more pronounced in the WT as compared with the *vir3-1* mutant (Fig. 6 **d**-**e**).

These experiments reaffirm stress regulation of VIR3 abundance, and the VIR3 role in reduced stability and/ or translational capabilities of GAPB and tAPX enzymes, and the formation of plastidial structure, stromules.

## Discussion

Dynamic regulation of plastids via complex and intertwined network of sensing, metabolic, and signaling functions guides’ cellular homeostasis, and shapes plant growth and development in an ever-changing environment. The data presented here reveal a multifaceted functional link between retrograde signaling and its signal transduction pathway component, VIR3. The finding specifically establishes stress-induced MEcPP-mediated increase in VIR3 levels, and an inverse correlation between VIR3 abundance and the levels of its interacting proteins, GAPB and tAPX, and the resulting shifts in plastidial metabolic and structural states.

Specifically, ample evidence obtained from a suppressor screen and the results from various loss-of-function and gain-of-function genotypes, together with those of inducible MEcPP producing lines (*HDSi* and *HDSi/vir3-1*) verifies VIR3 as a MEcPP signal transduction pathway component. Moreover, the significance of the zinc-binding domain in the VIR3 ability to complement the light-stress-induced variegation phenotype of *vir3-1* mutant corroborates the previous report using VIR3^H235L^ mutant construct (Qi *et al*., 2016). This data together with the identification of tAPX and GAPB as the VIR3 interacting proteins, and the inverse correlation between their levels and the VIR3 abundance provides support for the metalloprotease nature of this enzyme. It is of note that the positive correlation between stress-induced accumulation of MEcPP and increased abundance VIR3 is not supported by heightened transcript levels but by the enhanced translational capability and/or stability of VIR3 protein. This finding suggests a potential stress-induced MEcPP-mediated deactivation of a yet unknown protease that may target VIR3 under standard conditions.

Stress-induction of MEcPP-mediated increase in VIR3 abundance is ensued by reduced levels of tAPX, the enzyme responsible for detoxification of H_2_O_2_, a key signaling molecule with relatively long half-life that can alter cellular redox state and initiate Ca^2+^ signaling (Neuenschwander *et al*, 1995). In fact, Ca^2+^ signaling is necessary for induction of genes such as *ICS1*, whose expression is regulated by MEcPP-mediated induction of the *cis*-element *RSRE*, in a calcium dependent manner (Benn *et al*., 2016). These reports support the previously reported dose-dependent increase of SA in response to increasing H_2_O_2_ concentration (Neuenschwander *et al*., 1995), and further offer a potential molecular mechanism for reduced SA levels in *Rceh4* and *ceh1/vir3-1* versus the heightened SA levels in *ceh1/vir3-1/VIR3OE* lines (Fig. 1**d**). Another consequence of increased H_2_O_2_ is inhibition of the HDS enzyme activity and the enhanced levels of MEcPP (Wang *et al*., 2020).

Another outcome of stress-induced MEcPP-accumulation and heightened VIR3 levels is decreased levels of GAPB enzyme available for the conversion of G3P to **1**,3BPG, resulting in an increased G3P levels (Fig. 4**i**). This in turn could promote the MEP-pathway flux via escalating condensation of G3P with pyruvate, as the first step of the pathway (C. Obiol-Pardo, 2011; Lange *et al*., 2000; Tritsch *et al*., 2010; Zeng & Dehesh, 2021). This increase in flux, together with stress-induced reduced activity of the key enzyme, HDS (Li & Sharkey, 2013; Ostrovsky *et al*, 1998; Ostrovsky *et al*, 1992; Rivasseau *et al*, 2009; Walley *et al*., 2015; Wang *et al*., 2020; Wang *et al*., 2017; Xiao *et al*., 2012), not only support the consequential enhanced MEcPP levels, but also permits continuation of the flux through the pathway necessary for production of the essential metabolites, IPPs. The inverse correlation between GAPB abundance and MEcPP levels is best evident in comparative analyses of HL-treated WT versus *vir3-1* mutant plants. The analyses specifically illustrate higher abundance of GAPB and lower MEcPP levels in *vir3-1* compared to the WT plants (Figs. 6**a** and **c**). Accordingly, the continuation of the MEP-pathway flux in constitutively high or inducible MEcPP accumulating *ceh1/vir3-1* and *HDSi/vir3-1* lines is likely via compensatory routes other than the fine-tuning of GABP levels (Figs. 1**c** and 4**f**). One such a compensatory function could be via inhibition/reduction of GAPB enzyme activity by ROSs that oxidize the catalytic cysteine in the enzyme active site, a notion supported by the earlier reports concerning cytosolic GAPDH family members (Hancock *et al*, 2005; Zaffagnini *et al*., 2013). Moreover, stress-induced accumulation of ROS may in turn be regulated by GAPDHs, corroborating moonlighting function of these isozyme in response to biotic stresses (Henry *et al*., 2015).

The shift in H_2_O_2_, MEcPP and SA levels, is accompanied by formation of stromules, plastidial structures formed in response to range of stresses, and postulated to function as a conduit for transfer of signaling factors from chloroplast to nuclei (Caplan *et al*., 2015; Kumar *et al*, 2018). The differential prominence of these structural phenotypes in *ceh1* versus *ceh1/eds16*, (Figs 5**d**-**e** and supplemental Fig. 4**b**-**c**), and between HL-treated WT versus *vir3-1* mutant (Fig. 6**d**-**e**), are likely due to accumulation of both SA and H_2_O_2_ as the result of MEcPP-mediated enhanced abundance and/or stability of VIR3 and the consequential lowered levels of GAPB and tAPX enzymes.

The schematic model (Fig. 7) is a simplified depiction of the complex and intertwined reciprocity between MEcPP and VIR3 that provides a previously unrecognized link between the stress-induced plastidial retrograde signaling metabolite and a putative zinc-binding metalloprotease. This reciprocity provides insight into the molecular and biochemical venues involved in selective regulation of plastidial proteome and the consequential reconfiguration of the plastidial metabolic and structural states in response to environmental stimuli. This finding further signifies plastidial function in sensing multi-environmental inputs, integrating intraorganellar communication and ultimately mapping the cellular output central to maintenance of organismal integrity.

**Fig. 7.**
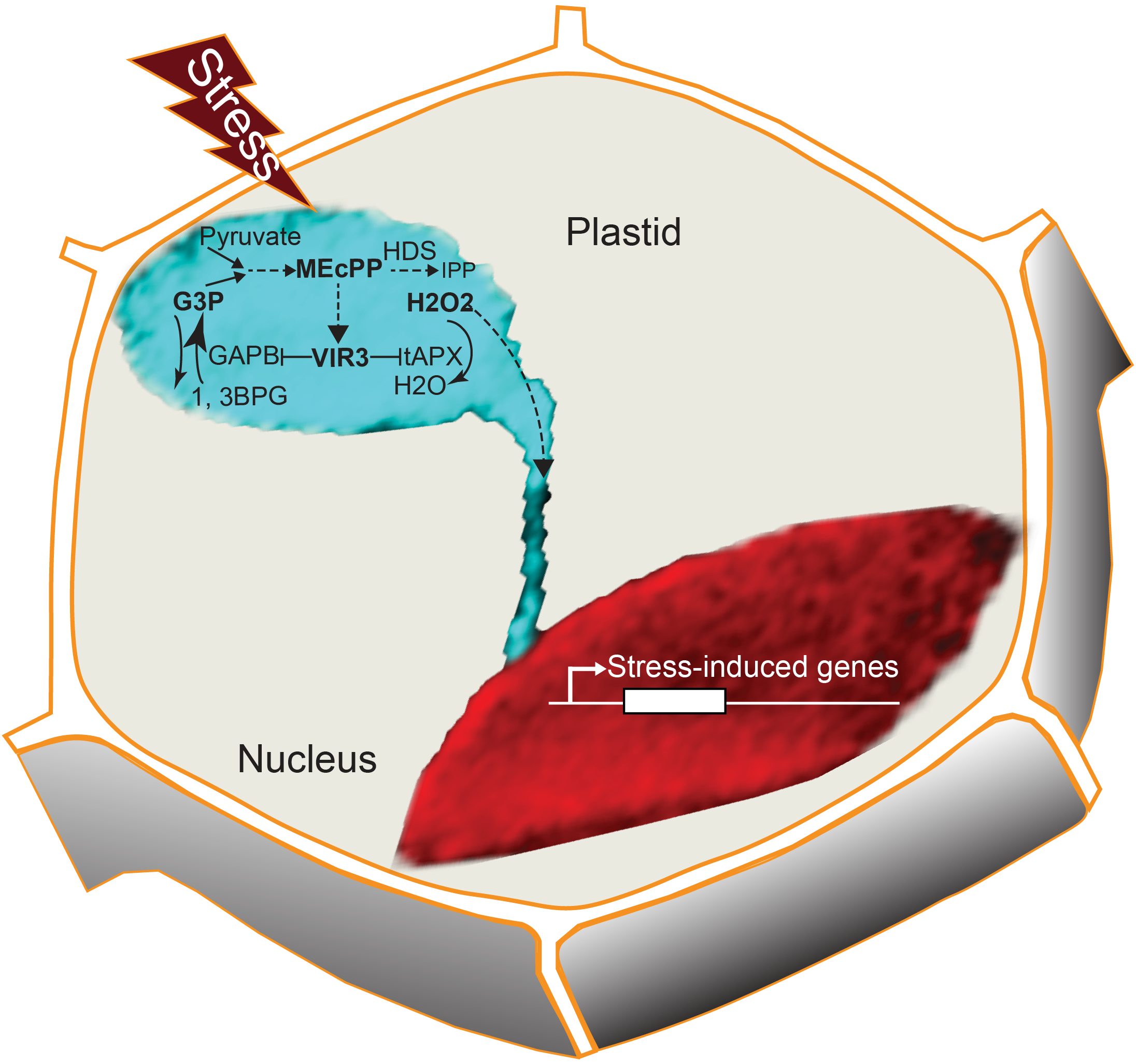
The schematic model depicting the reciprocity between MEcPP and VIR3 attuning the plastidial metabolic and structural states in response to environmental stimuli, thereby enabling retrograde regulation of selected stress-inducible nuclear genes.

## METHODS

### Plant materials

All Arabidopsis lines used in this research are in the background of Columbia-0. The *ceh1* mutant and DEX-inducible MEcPP accumulating line (*HDSi*) have been previously described (Jiang *et al*, 2020; Xiao *et al*., 2012). *Rceh4*, is the ethylmethylsulfonate (EMS) mutagenized obtained from the previously outlined screen (Jiang *et al*., 2020). The *vir3-1* and *p35S::VIR3-Flag/vir3-1* genotypes initially described (Qi *et al*., 2016), were provided by Dr. Yu (State Key Laboratory of Crop Stress Biology for Arid Areas and College of Life Sciences, Northwest Agriculture and Forestry University, Shaanxi 712100, People’s Republic of China) These lines were introgressed into *pHPL::LUC*, *ceh1*, and *HDSi*, and screened for homozygous lines. The *ceh1* and *ceh1*/*vir3-1* were also crossed with line expressing chloroplast CFP and mcherry-nuclear makers. Primers used for genotyping are listed (Supplemental Table 2)

Generation of *vir3-2* was based on CRISPR/Cas9 technology (Fauser *et al*, 2014). Specifically, we designed sgRNA (CCGTGGTGTGATATTGGATC) to target the 4th exon of *VIR3* (At1g56180) and cloned it into pEn-Chimera, and recombined with pDe-Cas9 using Gateway cloning. The construct was introduced into *ceh1* via *Agrobacterium* mediated transformation. The kanamycin selected transformants were sequenced over the 4th exon to select CRISPR lines containing deletions or insertions. The next generation of transformants were selected for the mutation and Cas9 segregation.

The *VIR3^H239Y^*-Flag construct was generated by site directed mutagenesis. Both *VIR3^H239Y^*-Flag and *VIR3*-Flag were amplified using *p35S::VIR3-Flag* as template and cloned into pENTR-D/Topo. All constructs were transferred to PUB-DEST vector to generate *pUBQ::VIR3^H239Y^*-Flag and *pUBQ::VIR3*-Flag constructs subsequently used to transform *vir3-1* line.

### Growth conditions

All seeds were surface-sterilized and sown on half strength Murashige and Skoog medium (2.16 g/L MS basal medium, 1 g/L MES (2-(N-morpholino) ethanesulfonic acid), pH 5.7, and 8 g/L phytagar). For selection tissue culture medium was supplemented with 50 mg/L kanamycin, 10 mg/L glufosinate ammonium (basta) or 25 mg/L hygromycin. After stratification for 2-3 days at 4°C in the dark, plates were transferred to a growth chamber at 22 °C with long day (16 h light and 8 h dark), and ~100 *μ* mol m^-2^ s^-1^ light intensity. After 14 days areal parts were harvested and immediately frozen in liquid nitrogen and stored at −80 °C until analysis.

High-light treatment was performed with plate grown seedlings on half strength MS at 22 °C with 16 h light/8 h dark, and exposed to high light (800-900 *μ* mol m^-2^ s^-1^ light intensity) for 2 days, followed by harvesting the areal parts after the treatment.

Plants were grown in constant light at ~100 *μ* mol m^-2^ s^-1^ light intensity.

*Nicotiana benthamiana* (tobacco) seeds were sown on Sunshine mix1 and grown for 6 weeks at 23 °C, and in long day condition and at ~100 *μ* mol m^-2^ s^-1^ light intensity.

### LUC split assay constructs

Upon amplification of *VIR3*, *VIR3^H239Y^*, tAPX (At1g77490) and GAPB (At1g42970), the fragments were cloned into pDONR207 and sequenced. The constructs were then transferred to vectors pMK7WG-cL-2, pMK7WG-nL-2, pMK7-cL-WG2 and pMK7-nL-WG2 using Gateway technology, resulting to generation of constructs: *p35S::VIR3-cLUC*, *p35S::VIR3-nLUC*, *p35S::cLUC-VIR3*, *p35S::nLUC-VIR3*, *p35S:: VIR3^H239Y^-cLUC*, *p35S:: VIR3^H239Y^-nLUC*, *p35S*::*cLUC-VIR3^H239Y^*, *p35S*::*nLUC-VIR3^H239Y^*, *p35S*::*tAPX-cLUC*, *p35S*:: *tAPX-nLUC*, *p35S*::*cLUC*-*tAPX*, *p35S*::*nLUC-tAPX*, *p35S*::*GAPB-cLUC*, *p35S*:: *GAPB-nLUC*, *p35S*::*cLUC-GAPB*, *p35S*::*nLUC-GAPB*, *p35S*::*cLUC*, *p35S*::*nLUC*.

### Marker lines generation and confocal microscopy

Nuclear markers were generated by amplification of the WPP domain of RANGap1 from pGLP2::NTF and the mcherry from pBL14, flanked by the ACT2 promoter and 35S terminator amplified from pACT2::BirA. These four fragments and the pMDC123 vector linearized by SpeI and AscI were assembled together by NEB Gibson assembly. Plastidial marker was generated by amplification of genomic DNA from pt-yk transgenic plants containing rubisco plastidial transit peptide fused to CFP cassette (pt-CFP) (Nelson *et al*, 2007). Next, the two cassettes were assembled into pMDC123 vector by NEB Gibson Assembly to generate the nucleus-mcherry-pt-CFP marker construct, subsequently employed for transforming into the *ceh1* background, and eventually transgressed into other genotypes. These genotypes were employed in confocal microscopy imaging. Additionally, we employed RB-GFP and NRIP1-Cerulean expressing lines, the generous gifts from Professors Steve Theg and Savithramma Dinesh-Kumar at UC Davis.

An upright confocal microscope (Zeiss LSM 880 Upright Laser Scanning Confocal Microscopy) equipped with 40X/1.2 water or 40X/1.4 Oil objectives was used for fluorescent protein marker detection.

### MEcPP and SA measurements

Extraction and quantifications of MEcPP and SA were performed as previously described (Jiang *et al*., 2020; Jiang *et al*., 2019).

### Luciferase measurements

Fourteen days old plants grown on half strength MS were sprayed with 1 mM Luciferin (Gold Biotechnology, LUCK-1G) with 0.001% triton X-100, and immediately imaged 10x for 15 min each time using Andor DU434-BV CCD camera (Andor Technology). Luciferase activity was quantified as described before (Walley *et al*, 2007; Wang *et al*, 2014).

### Tobacco infiltration

Tobacco leaves of 6 wk. old plants were infiltrated as described previously(Wang *et al*., 2014). Cultures of constructs grown in *Agrobacterium tumefaciens* GV3101 were prepared in infiltration medium (50 mM MES, 9.8 mM MgCL_2_-6H_2_O, 27.7 mM D-glucose, 0.1 mM acetosyringone). Before infiltration, cultures were diluted to OD_600_ of 0.1 and mixed in equal amounts. P19 was infiltrated with all samples.

### Immunoprecipitation assay

Immunoprecipitation (IP) was performed 72 hr after infiltration of tobacco leaves followed by harvesting of these leaves immediately frozen in liquid nitrogen. Proteins were extracted in extraction buffer (10 % glycerol, 25 mM Tris pH 7.5, 150 mM NaCl, 2 % polyvinylpolypyrrolidone, 10 mM dithiothreitol, 1% proteinase inhibitor without EDTA), centrifuged for 10 min at 14000 rpm, and the supernatant was incubated with 10 *μ*l packed anti-Flag M2 beads (Millipore-Sigma) for 4 hr, at 4°C. Beads were washed 4x with wash buffer (10 % glycerol, 25 mM Tris pH 7.5, 150 mM NaCl, 2 % polyvinylpolypyrrolidone, 10 mM dithiothreitol) and 1x with 1 mL of ammonium bicarbonate solution (50 mM, pH 8). On-beads trypsin digestion was performed as previously described (Drakakaki *et al*, 2012).

IP assays using Arabidopsis lines were performed on proteins obtained from enriched chloroplasts as previously described (Shi *et al*, 2012). In short, areal tissue of 14 days old plants grown on half strength MS were homogenized in extraction buffer (1.25 M NaCl, 0.25 M ascorbic acid, 10mM sodium metabisulfite, 0.0125 M sodium tetraborate, 50mM Tris pH 8.0, 1 % PVP-40, 0.1 % BSA, 1 mM dithiothreitol, pH 3.8). Homogenate was filtered through Miracloth and centrifuged at 200 g for 20min, followed by centrifugation of supernatant at 200 g for 20 min, and thereafter at 3.500 g for 20 min. Pellet was washed with wash buffer (1.25 M NaCl, 0.0125 M sodium tetraborate, 50mM Tris pH 8.0, 1 % PVP-40, 0.1 % BSA, 1 mM dithiothreitol, pH 8.0) and centrifuged at 3750 g for 20 min. Pellet was re-suspended in IP buffer (1x PBS, 0,3 % triton X-100, 1%, 1% proteinase inhibitor without EDTA) and mixed with anti-Flag M2 beads and continued described above. After the 4th wash, beads were mixed with 2x SDS loading buffer and used in immunoblot analyses.

### Protein extraction and immunoblot analysis

Soluble and membrane bound Proteins were extracted from 14 day old areal tissue according to the previously described method(Gou *et al*, 2015). Soluble protein were separated on a 12% SDS-page gel and membrane protein were loaded on a 15% Urea-SDS page gel as described (Nakatogawa *et al*, 2012)and subsequently transferred onto PVDF membranes. Membranes with soluble protein were probed with anti-GAPB (gift from Dr Zeeman, ETH Zurich, Switzerland) (1:3000), whereas membrane protein blots were probed with anti-APX (Agrisera, AS08 368**) (**1:2.000), and with anti-Flag M2 (Millipore-Sigma) (1:1000) or anti-αTIP (ABRC, AB00111) (1:1000). Accordingly, secondary antibodies selected included anti-rabbit-HRP (ThermoFisher, 31460) (1:20.000), anti-mouse-HRP (KPL, 074-1806) (1:10.000), anti-chicken-HRP (KPL, 14-24-06) (1:10.000) or anti-chicken-phosphatase (KPL, 15-24-06) (1:3000). Visualization was achieved using chemiluminescent reaction substrate (ThermoFisher, supersignal west pico plus substrate) according to manufacturer’s instructions and ChemiDoc PM (Bio-Rad).

Protein band intensities were measured and normalized using Image Lab 6.0.1 software (BioRad).

### Proteomics

2D-nano LC/MS analysis of trypsin digested proteins was performed as previously described (Drakakaki *et al*.,2012; Zhu *et al*, 2018). Proteome Discoverer 2.1 (Thermo Scientific) was used to analyze the raw MS data files. MS data were matched to Niben1.0.1 and NbS tobacco, TAIR10 Arabidopsis, and SL4.0 tomato databases (Supplemental Table 1).

### Quantitative gene expression analyses

Total RNA was extracted and used for RT-qPCR as previously described (Wang *et al*., 2017). The normalization was carried out using the internal controls AT4G26410. Each experiment was performed on three biological replicates each with three technical replicates. Primers used to detect VIR3, tAPX and GAPB were listed in Supplementary Data Table 2qPCR.

### Statistical analyses

All experiments were performed with at least three biological replicates. Data are mean ± standard deviation (SD). The analyses were carried out via a two-tailed Student’s *t* tests with a significance of *P* < 0.05.

The primers used in the construction are summarized in the Supplemental Table 2.

## Data availability

Proteome database of VIR3 binding proteins is provided in Supplemental Table1. Any additional data or biological material that support the findings of this study are available from the corresponding author upon reasonable request.

## Author Contributions

J.W., W.V., Y.X. and K.D. designed the study, and Y.X. performed the mutagenesis and isolated the *Rceh1* mutant, J.W., W.V., X.H., H.K., P.Y. and A.S. performed the experiments, and K.D. wrote the manuscript.

## Acknowledgements

We would like to thanks Prof. Fei Yu (Northwest Agriculture and Forestry University) for providing vir3-*1* and *p35S::VIR3-Flag/vir3-1* genotypes, and Prof Samuel Zeeman (Institute of Molecular Plant Biology, ETH Zurich, Switzerland) for proving GAPB specific antibody. We would like to extend our gratitude to Professors Steve Theg and Savithramma Dinesh-Kumar at UC Davis for providing lines expressing RB-GFP and NRIP1-Cerulean, respectively. This work was funded by Kathy Cookson and John Leibacher Endowed Chair fund, and by National Institutes of Health (NIH) R01GM107311 to KD.

